# Whole blood transcriptome analysis in bipolar disorder reveals strong lithium effect Authors

**DOI:** 10.1101/497784

**Authors:** Catharine E. Krebs, Anil P.S. Ori, Annabel Vreeker, Timothy Wu, Rita M. Cantor, Marco P. Boks, Rene S. Kahn, Loes M. Olde Loohuis, Roel A. Ophoff

## Abstract

Bipolar disorder (BD) is a highly heritable mood disorder with complex genetic architecture and poorly understood etiology. We performed a whole blood transcriptome analysis in a BD case-control sample (*N*_subjects_ = 480) by RNA sequencing. While we observed widespread differential gene expression patterns between affected and unaffected individuals, these effects were largely linked to lithium treatment at the time of blood draw (FDR < 0.05, *N*_genes_ = 976) rather than BD diagnosis itself (FDR < 0.05, *N*_genes_ = 6). These lithium-associated genes were enriched for cell signaling and immune response functional annotations, among others, and were associated with neutrophil cell-type proportions, which were elevated in lithium users. Neither genes with altered expression in cases nor in lithium users were enriched for BD, schizophrenia, and depression genetic risk based on information from genome-wide association studies, nor was gene expression associated with polygenic risk scores for BD. Our findings suggest that BD is associated with minimal changes in whole blood gene expression independent of medication use but underline the importance of accounting for medication use and cell type heterogeneity in psychiatric transcriptomic studies. The results of our study add to mounting evidence of lithium’s cell signaling and immune-related mechanisms.

## Introduction

Bipolar disorder (BD) is a chronic and recurrent psychiatric disorder affecting approximately 1% of the population worldwide and presenting a major public health burden^1, 2^. It is characterized clinically by instability in mood resulting in manic and depressive episodes interspersed between neutral, euthymic states^2^. Risk for BD is highly genetic, with heritability estimates as high as 85%^3^ and common genetic variation explaining up to a third^4^. Still, however, the pathophysiological characteristics of BD are not well understood. Investigating molecular phenotypes such as gene expression as intermediate measures between genetic variation and clinical variation is a viable strategy for uncovering disease mechanisms. Many such studies have been carried out for BD, and in Table 1 we present a summary that reveals a lack of consistency between findings likely owing to clinical heterogeneity, differing study designs, and the low numbers of samples investigated (*N* ≤ 62 BD subjects)^5–27^. Moreover, there are many potential confounds that impact gene expression, including medication.

**Table 1.**
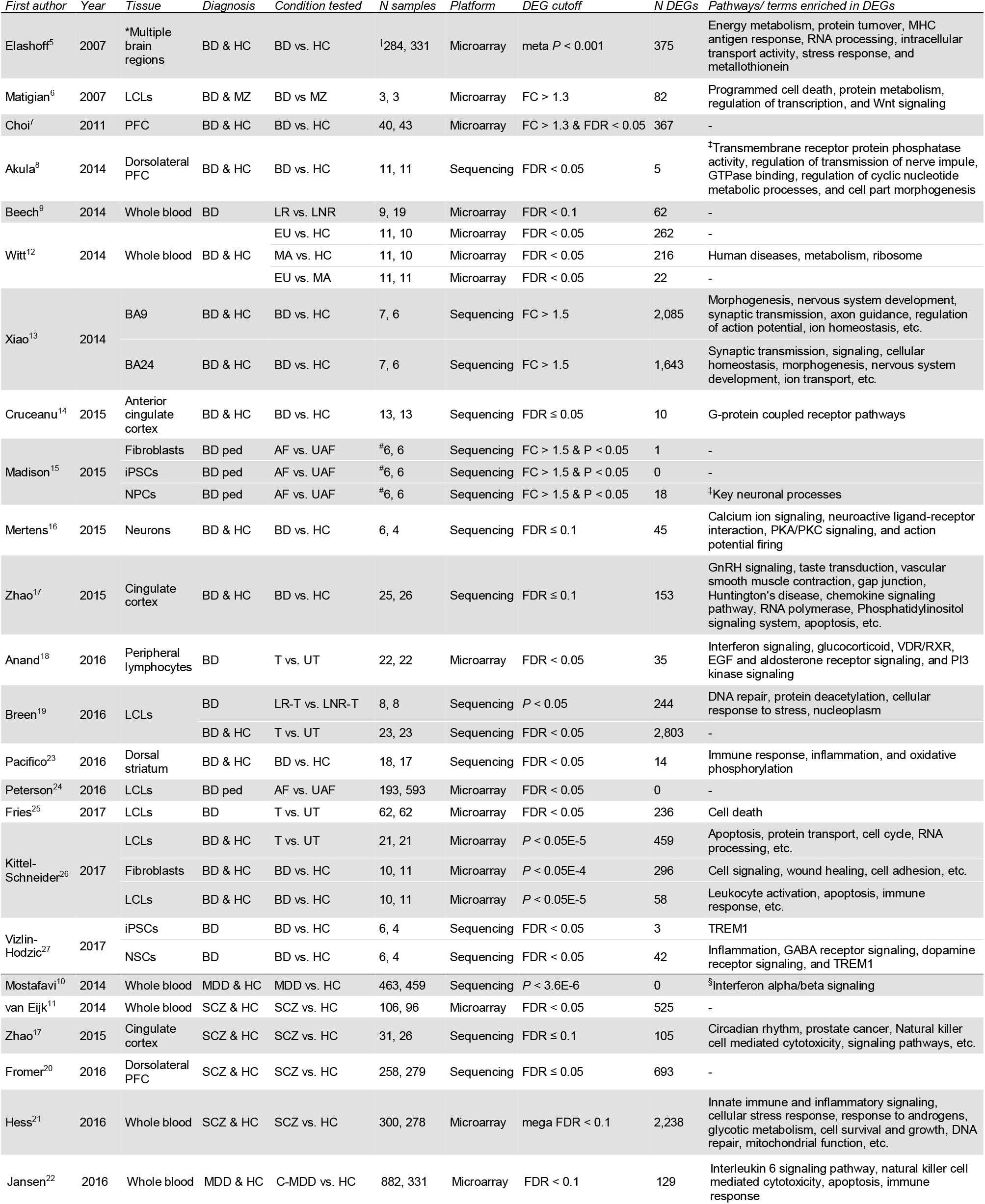
Review of previous BD and lithium studies with differential expression analyses. Select schizophrenia and major depressive disorder studies were included (at the bottom of the table) as examples of what larger BD and lithium studies might look like. *Multiple brain regions including frontal BA46, BA10, BA6, BA8, BA9, and cerebellum. ^†^165 BD individuals (samples partially overlapping). ^‡^Enrichment analysis was performed on genes with nominal p-values (P < 0.05). §Enrichment analysis was performed on genes with small p-values (sets of top *N* genes, *N* = [30, 60, 100, 150, 300, 500]). #N = 2 samples with 3 replicates each. Abbreviations: AF, affected; BD, bipolar disorder; BD ped, BD pedigree; C-MDD, current major depressive disorder; DEGs, differentially expressed genes; EU, euthymic; FC, fold change; FDR, false discovery rate; HC, healthy control; iPSCs, induced pluripotent stem cells; LCLs, lymphoblastoid cell lines; LNR, lithium non-responder; LNR-T, lithium non-responder treated with lithium; LR, lithium-responder; LR-T, lithium responder treated with lithium; MA, manic; MDD, major depressive disorder; MZ, unaffected monozygotic twin; NPCs, neural progenitor cells; NSCS, neural stem cells; PFC, prefrontal cortex; SCZ, schizophrenia; T, treated with lithium; UAF, unaffected; UT, untreated with lithium.

Therefore, to explore gene expression changes associated with BD, we generated RNA sequencing data from peripheral whole blood collected in a large, well-characterized case-control cohort from The Netherlands. We examined gene expression differences between groups both at the individual gene level and at the level of gene co-expression. Upon correction for technical and biological variables including the use of lithium, the most widely used prescription drug in our cohort, gene expression differences between subjects with BD and controls were minor. Differences in subjects being treated with lithium compared to those who are not, however, were widespread. These differences were partially but not entirely explained by differences in cell-type composition, driven by elevated neutrophil proportions in lithium users. The lithium-associated changes in gene expression were independent of psychiatric genetic risk, though. Our results suggest nominal BD-related gene expression effects in blood but numerous effects related to lithium treatment. This work highlights the importance of accounting for medication use in psychiatric transcriptomic studies and provides insight into lithium’s molecular mechanisms of action.

## Methods

### Sample preparation and RNA sequencing

See Supplementary Methods for more information regarding sample ascertainment and assessment. Peripheral whole blood was drawn and processed for genotyping and RNA sequencing from 240 controls and 240 cases, of whom 227 (94.6%) had a diagnosis of bipolar I disorder and 13 (5.4%) had a diagnosis of bipolar II disorder. Whole blood was collected in PaxGene Blood RNA tubes and total RNA extracted using the PAXgene isolation kit (Qiagen) according to manufacturer’s protocols. RNA integrity number (RIN) values were obtained using Agilent’s NRA 6000 Nano kit and 2100 Bioanalyzer. RNA concentrations were determined using the Quant-iT RiboGreen RNA Assay kit. The UCLA Neuroscience Genomics Core subsequently performed RNA sequencing and prepared sample libraries using the TruSeq Stranded RNA plus Ribo-Zero Gold library prep kit to remove ribosomal and globin RNA to enrich for messenger and noncoding RNAs. Concentration of the sequencing library was determined on a TapeStation and a pool of barcoded libraries were layered on eight lanes of the Illumina flow cell bridge amplified to raw clusters. An average of 24.9 million paired-end reads of 75 bases in length per sample were obtained on an Illumina HiSeq 2500. The raw sequence data were processed for quality control (QC) using FastQC, after which all samples were deemed suitable for downstream analysis.

### RNA sequencing alignment and gene expression quantification

Reads were mapped to human reference genome hg19 using TopHat2^28^ allowing for two mismatches yielding an average mapping rate of 96.0% per sample and an average concordant pair mapping rate of 89.8% per sample. Samples had an average of 33.9% duplicate reads. Picard Tools were used to obtain 18 different sequencing metrics such as number of reads, percent mapped reads, and number of coding bases, that were examined for QC and then processed for dimension reduction using principal component analysis (PCA; Supplementary Methods). The first three principal components, which explain 75.9%, 16.9%, and 6.4% of variance, respectively, were used as covariates in subsequent analyses. Known Ensembl gene levels were quantified using HTSeq in the union mode to obtain integral counts of reads that intersect the union of all transcripts of genes. PCA of gene expression quantification was used for data visualization and additional QC, after which four samples were removed for apparent mix-up (Supplementary Methods). Thirty-two additional samples were excluded due to missing demographic information. Differential expression and co-expression analyses were therefore limited to a set of 444 subjects (240 cases and 204 controls).

### Normalization, covariate correction, and differential expression analysis

Gene expression counts from HTSeq were filtered for genes having > 10 counts in 90% of samples, yielding 12,344 genes for subsequent analyses. Filtered counts were converted to log2-counts-per-million (log-cpm) to account for differences between samples in sequencing depth and to stabilize variances at high counts. Then, the mean-variance relationship was modelled with precision weights at the individual observation level using limma voom^29^. Briefly, voom non-parametrically estimates the mean-variance trend of the logged read counts and uses this to predict the variance of each log-cpm value. The predicted variance is then used as a weight, which is incorporated into the linear model procedure during differential expression analysis. These gene-wise weighted least-squares linear models are fitted to the normalized log-cpm values, taking into account the voom precision weights and the final covariate model, generating a coefficient for the effect of each variable on each gene’s expression:

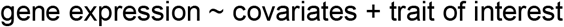

Then, for each gene, the coefficient for the trait of interest is statistically tested for being significantly different from zero. *P*-values from this test were corrected for multiple testing using the Benjamini-Hochberg false discovery rate (FDR) estimation, and a gene was considered to be differentially expressed if it had an FDR-corrected *P*-value < 0.05. The final covariate model for differentially expressed genes (DEGs) between BD cases and controls included the following variables: age, sex, lithium use, tobacco use, assessment group, RIN, sequencing plate, and sequencing metric PCs 1 through 3. The final covariate model for DEGs between subjects being treated with lithium (i.e. lithium users) and non-lithium users included the following variables: BD diagnosis, age, sex, tobacco use, assessment group, RIN, sequencing plate, and sequencing metric PCs 1 through 3. Tobacco use was included because of its well-characterized effect on whole blood gene expression^30^. An overview of covariates can be found in Table S1. DEGs were checked for overlap and concordance with other datasets (Supplementary Methods). Fold changes (FC) reported are in log2 fold change units.

### Co-expression network analysis

To determine networks of genes with correlated expression, weighted gene co-expression network analysis (WGCNA)^31^ was performed using the WGCNA package in R. To do this, first the 12,344 filtered and normalized genes were residualized adjusting for the following covariates: age, sex, tobacco use, assessment group, RIN, sequencing plate, and sequencing metric PCs 1 through 3. Then, briefly, WGCNA defines a network of genes as nodes with edges between genes based on pairwise correlations between genes, and separates the network into modules of gene clusters with highly coordinated expression. The *β* parameter (β = 7) was chosen according to the approximate scale-free topology criterion described by Langfelder and Horvath^31^. Then the gene expression profiles of each module were summarized by calculating the module eigengene, which is defined as the first principal component of the expression matrix of that module. Each gene was then assigned a measure of module membership for each module.

To determine biologically significant modules, gene significance measures were assigned to each gene for each of our traits of interest, including BD diagnosis and lithium use, by calculating the absolute correlation between the trait and the expression profiles. Then a measure of module-trait significance was calculated by correlating module membership values with gene significance values. An association was considered significant if its P-value surpassed Bonferroni correction for testing multiple modules (*P* < *α* = 0.05/*N*_modules_). Finally, intramodular connectivity *k*_IM_ was calculated to determine the level of connectivity for the genes in modules significantly associated with traits of interest.

### Functional annotation

The Database for Annotation, Visualization, and Integrated Discovery (DAVID, v6.8)^32^ was used for functional annotation of each gene list. We used three gene lists from the differential expression analysis: the 976 lithium DEGs at FDR < 0.05, the 754 up-regulated lithium DEGs at FDR < 0.05, and the 222 down-regulated lithium DEGs at FDR < 0.05. We also used gene lists from the five co-expression network analysis modules that were significantly associated with BD: M1 (*N*_genes_ = 2,092), M7 (*N*_genes_ = 700), M9 (*N*_genes_ = 55), M11 (*N*_genes_ = 622), and M26 (*N*_genes_ = 484). The full set of 12,344 filtered and normalized genes used as input for differential expression and co-expression network analyses was used as background to determine overrepresentation in each of the gene lists. The functional annotation clustering tool was applied using unique Ensembl IDs and the following databases: SP_PIR_KEYWORDS, UP_SEQ_FEATURE, GOTERM_BP_FAT, GOTERM_CC_FAT, GOTERM_MF_FAT, BIOCARTA, KEGG_PATHWAY, INTERPRO, UCSC_TFBS. Cluster annotations were called significant if the enrichment was greater than 1.0 and at least 1 gene list in the annotation cluster survived Bonferroni correction (*P* < 0.05).

### Estimation of cell-type proportions

To estimate cell-type composition in our sample we employed the CIBERSORT online software (cibersort.stanford.edu)^33^. Briefly, CIBERSORT uses reference gene expression signatures to estimate the relative proportions of cell types in tissues with complex, heterogeneous cell composition via linear support vector regression. The reference dataset we used to deconvolve our mixture of whole blood cell types was the validated leukocyte gene signature matrix that is provided with the CIBERSORT software, termed LM22^33^. It contains 547 genes whose expression discriminate between 22 different human hematopoietic cell phenotypes (Table S2), including seven T-cell types, naive and memory B cells, plasma cells, natural killer cells, and myeloid subsets.

To prepare our gene expression data for input to CIBERSORT, raw expression counts from HTSeq were converted to transcripts per million (TPM). Using the resulting matrix of TPM values for our 480 samples and the LM22 gene signature matrix as input, CIBERSORT was run online with 100 permutations and with quantile normalization disabled as recommended for RNA-seq data. The output matrix consisted of deconvolution results with relative fractions of cell types normalized to 1 across all cell subsets for each sample. These estimated cell-type proportions were then residualized using a linear regression model adjusting for the following covariates: sex, age, tobacco use, sequencing plate, RIN, and sequencing metric PCs 1 through 3. Then, residualized cell-type estimates were used to predict lithium use in a stepwise linear regression using the stepAIC function in the MASS package in R. The estimated cell-type proportions were also appended to the table of technical and biological covariates and then used to re-run the differential expression analysis while accounting for cell-type heterogeneity in the sample.

### Enrichment of cell types in co-expression modules

The enrichment of LM22 cell types in gene co-expression modules determined from WGCNA was calculated in two ways. First, the hypergeometric overlap between modules and cell type signature genes was calculated. The binary matrix of LM22 signature genes provided by Newman et al.^33^, where 1 denotes that a gene was significantly differentially expressed in that particular cell type and 0 denotes that it was not, was used to extract lists of signature genes for each cell type, or genes with a value of 1. These lists are partially overlapping, with 262 genes being unique to a given list and 285 genes being shared between ≥ 2 lists (maximum 10 lists). Then, using the GeneOverlap library in R, the hypergeometric overlap was calculated between each of these 22 cell type signature gene lists and each of the 27 module gene lists using the full set of 12,344 filtered and normalized genes as background.

Second, binary cell type signatures were used to predict module membership values in a linear model. We reasoned that this method might be more powerful than a strict overlap due to the fact that every gene has a module membership value for every module, regardless if it was assigned to that module. The gene co-expression network output, which consists of module membership values for each gene for each module, was limited to the set of LM22 signature genes that were expressed in our sample (*N*_genes_ = 331). These values were then used as an outcome in a linear model, with the binary matrix of LM22 signature genes as predictors. To avoid multiple testing penalties, only five regressions were run on the five modules that were associated with lithium: M1, M7, M9, M11, and M26.

### Integration of GWAS data with transcriptomic signatures

Prior to gene-set analyses, heritability and genetic correlation of traits of interest were estimated to confirm significant non-zero SNP-based heritability (Supplementary Methods). Analyses were performed across three psychiatric genome-wide association study (GWAS) traits from publicly available datasets (bipolar disorder, schizophrenia, and self-reported depression) and 2 sets of DEGs (BD at FDR < 0.2 and lithium-use at FDR < 0.05). Differential expression log2 fold changes and FDR-corrected P-values for each of the 12,344 genes expressed at > 10 counts in 90% of samples were obtained from limma to integrate whole-blood gene expression signatures with GWAS data using Multi-marker Analysis of GenoMic Annotation (MAGMA v1.06)^34^.

GWAS summary statistics were obtained for the following three GWAS traits:

1. SCZ^35^: 36,989 cases and 113,075 controls;
2. BD^36^: 20,352 cases and 31,358 controls;
3. 23andMe self-reported depression^37^: 75,607 cases and 231,747 controls;

The 1000 Genomes Project Phase 3 release European reference panel (*N* = 503) was used to model LD in all analyses^38^. Eight gene lists were used from two different DEG models along with a positive and negative control:

1. Lithium-use DEGs at FDR < 0.05: *N* = 897 genes;
2. Up-regulated lithium-use DEGs at FDR < 0.05: *N* = 680 genes;
3. Down-regulated lithium-use DEGs at FDR < 0.05: *N* = 217 genes;
4. BD DEGs at FDR < 0.2: *N* = 630 genes;
5. Up-regulated BD DEGs at FDR < 0.2: *N* = 389 genes;
6. Down-regulated BD DEGs at FDR < 0.2: *N* = 241 genes;
7. Positive control gene-set: the top 100 most significant genes from a random draw of *N* = 1,000 using the BD GWAS gene-level test statistics;
8. Negative control gene-set: a random draw of *N* = 1,000 genes using the BD GWAS gene-level test-statistics.

MAGMA was used to run *gene property* analyses, which uses a multiple regression framework to associate a continuous gene variable to GWAS gene level p-values. High quality SNPs (INFO > 0.9) were mapped to genes using Ensembl gene IDs and NCBI build 37.3 gene boundaries +/- 10kb extensions using the -- annotate flag. For each phenotype, we generated gene-level p-values by computing the mean SNP association using the default gene model (‘snp - wise=mean’). We only included SNP with MAF > 5% and dropped synonymous or duplicate SNPs after the first entry (‘synonym-dup=drop-dup’). For each annotation, we then regressed gene-level GWAS test statistics on the corresponding gene annotation variable using the ‘--gene-covar’ function while adjusting for gene size, SNP density, and LD-induced correlations (‘--model correct=all’), which is estimated from an ancestry-matched 1KG reference panel. In all analyses, we included only genes for which we had both the gene variable and GWAS gene level test statistic available. Two-sided p-values are reported.

Secondary gene-set analyses were run on a limited number of DEG gene sets and additional, sleep-related GWAS traits (Supplementary Methods).

## Results

### Minimal changes in bipolar disorder gene expression

To explore the transcriptomic signatures of BD, we first evaluated whether subjects with BD harbored transcriptional differences on a per gene level compared with controls. Of the 12,344 genes tested, only six were differentially expressed in BD after correcting for multiple testing (FDR < 0.05; Figure 1A). The differences in expression were very small, with absolute fold changes ranging from 0.12 to 0.44. While the number of identified differentially expressed genes (DEGs) was too small to perform functional enrichment analysis, we did find that three of the six genes (*COG4, DOCK3*, and *BBS9)* were expressed in GTEx frontal cortex tissue (median TPM > 1) and show relatively stable expression across brain cell types except for *DOCK3*, which is enriched in neurons (fold change relative to other cell types = 6.82; Table S3). Four of the genes were present in the Stanley Genomics brain gene expression database, and two of these were found to be differentially expressed in BD individuals in at least one study, *COG4* and *DOCK3*, although the latter was altered in the opposite direction. *COG4* was also reported as differentially expressed in a schizophrenia mega-analysis of nine whole blood microarray datasets^21^. Using polygenic risk scores (PRS) for BD as the differential expression trait of interest rather than the dichotomous case-control phenotype did not yield any significant genes, even though PRS did significantly differ between BD cases and controls (*t* = −3.42, *P* = 6.88 x 10^-4^; Figure S1; Supplementary Methods).

**Figure 1.**
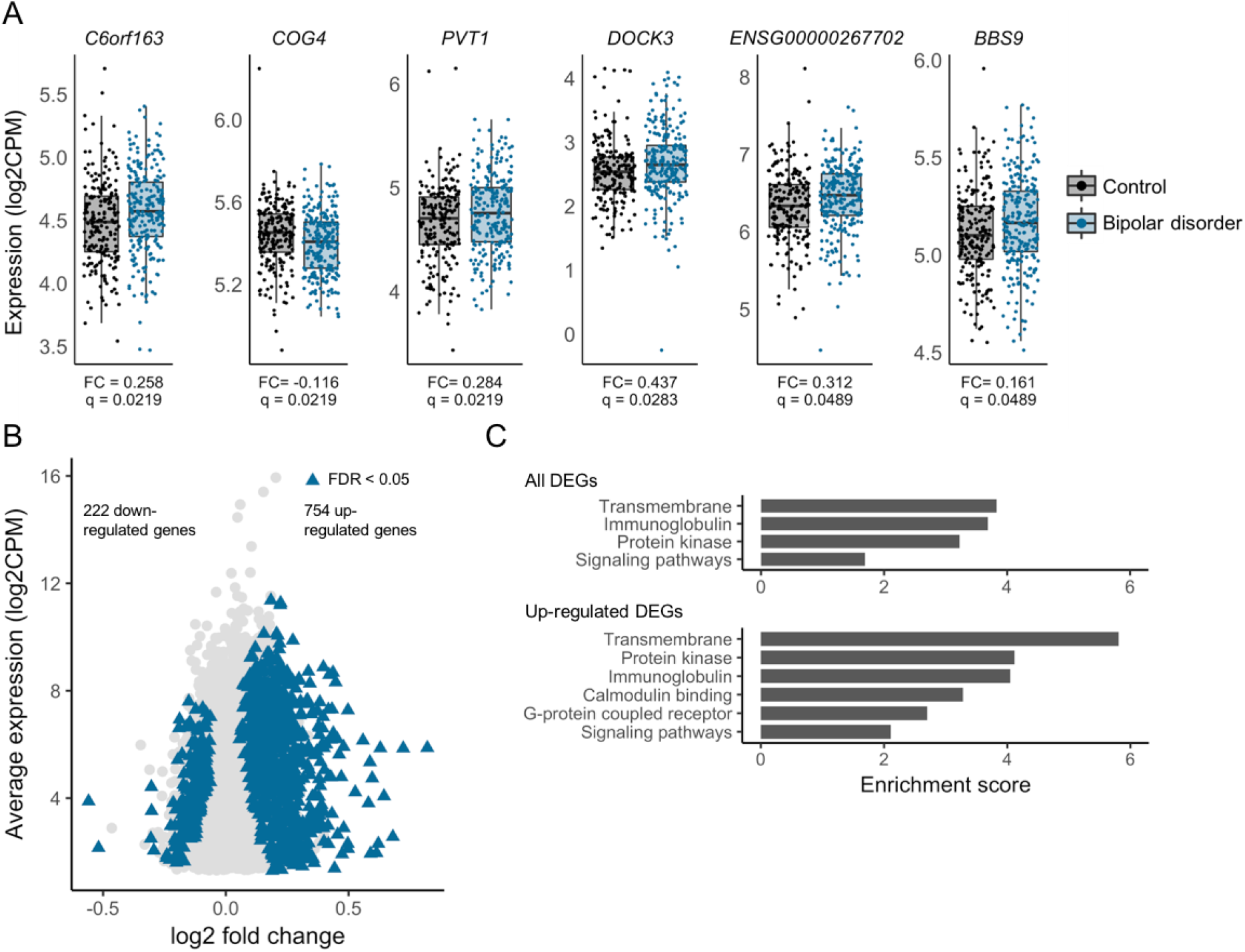
Differentially expressed genes. (A) Six BD DEGs. FC, log 2 fold change; q, FDR-adjusted *P* < 0.05. (B) 976 genes differentially expressed between lithium users and non-lithium users (shown as blue triangles, FDR-adjusted *P* < 0.05; all other genes tested shown as light gray circles). (C) DAVID^32^ functional annotation cluster enrichment of all 976 DEGs (upper) and 754 up-regulated DEGs (lower). Enrichment scores increase when the gene list is limited to up-regulated genes only. Clusters were considered significant if the enrichment score > 1 and at least one term in the cluster survived Bonferroni correction for multiple testing.

### Widespread subtle gene expression changes in lithium users

Following the same differential expression pipeline as above, we found 976 genes with small differences in gene expression between lithium users and non-lithium users (|FC| mean = 0.20, max = 0.82, SD = 0.10; Figure 1B, Supplementary File 1). These genes were enriched for biological terms related to calcium signaling and other signaling pathways, and immunity (Figure 1C). To distinguish between up- and down-regulated gene pathways, we stratified genes by their direction of change in expression. The 754 up-regulated genes were annotated for many of the same terms as the full set but with greater enrichment scores, indicating that the up-regulated genes are driving the enrichment scores in the full set (Figure 1C). Of the 976 lithium-use DEGs, 804 were expressed in GTEx frontal cortex samples (TPM > 1), and 488, 553, 503, 478, 512, and 403 were expressed in neurons, fetal astrocytes, mature astrocytes, oligodendrocytes, microglia/ macrophages, and endothelia, respectively (FPKM > 1). However, none of these gene sets were significantly enriched (hypergeometric P > 0.05).

To validate our results, the 976 lithium-use DEGs were tested for overlap with lists of DEGs from similar studies found in the literature (Table S4). Although none of these studies has the same design as ours, we did find a significant overlap between our 976 lithium-use DEGs and the lists from two studies. In the first study^18^, DEGs were detected by comparing peripheral monocyte gene expression in subjects before and after lithium monotherapy. Of the 35 DEGs discovered, 18 were shared with the current study (hypergeometric odd ratio (OR) = 13.57, *P* = 4.66 x 10^-12^), and all 18 were concordant in direction (Figure S2A). In the second study^19^, DEGs were detected by comparing LCL gene expression before and after lithium treatment *in vitro.* Of the 1,504 DEGs discovered, 134 were shared with our study (hypergeometric OR = 1.27, *P* = 9.23 x 10^-3^), and 84.6% of these were concordant in direction (Figure S2B). There were two genes shared between all three lists, *RFX2* and *SLC29A1.* We report genes in these overlapping lists as high confidence lithium-associated genes (Supplementary File 1).

### Modules of co-expressed genes are associated with lithium use

Next, in search of genes with differential co-expression in BD, we constructed a gene expression network in the entire sample using WGCNA and assessed the detected modules for association with BD. This network consisted of 27 modules ranging in size from 48 to 2,760 genes (mean *N*_genes_ = 441, Supplementary File 2). By evaluating the correlation of module membership values with gene significance for BD diagnosis, we quantified the association of each module with BD. After Bonferroni multiple testing correction, five modules were significantly associated with lithium-use, but no modules were associated with BD or any other clinical or technical variable (Table S5). Of the five modules associated with lithium use, three shared significant overlap with lithium-use DEGs (Table 2). M26 was most significantly associated with lithium (*P* = 2.00 x 10^-4^; Figure S3A) but was not significantly enriched for lithium DEGs. M1 was also associated with lithium (*P* = 9.04 x 10^-4^; Figure S3B) and had the most significant enrichment of DEGs (431 of 2,092 genes in the module were DEGs; hypergeometric OR = 4.62, *P* = 2.03 x 10^-97^). Functional annotation clustering of the genes in M1 showed an enrichment of terms related to cell signaling, immunity, and glycophosphatidylonositol anchor.

**Table 2.**
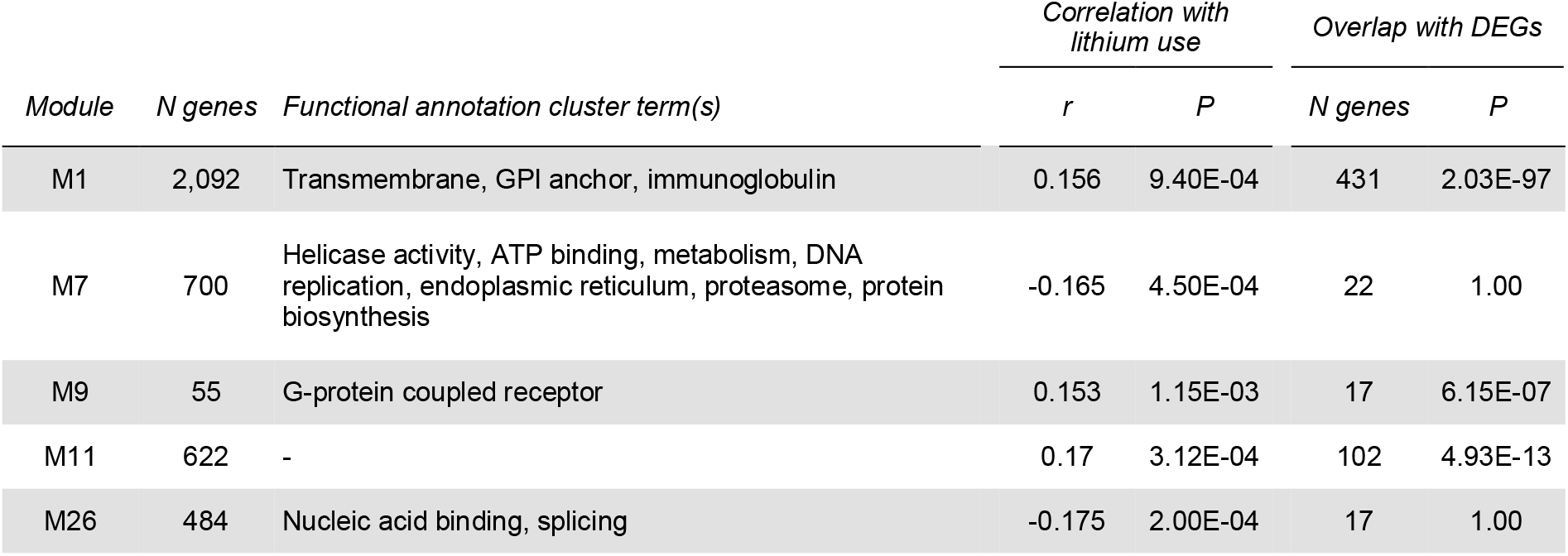
Co-expression module association with lithium use. Functional annotation cluster enrichment determined using DAVID^32^. Correlation with lithium use calculated by correlating gene module membership values with gene significance values for lithium use. Overlap was calculated by testing for hypergeometric overlap between the list of lithium-use DEGs and the list of genes within each module. Abbreviations: DEGs, differentially expressed genes; GPI, glycophosphaditylinositol.

| Module | N genes | Functional annotation cluster term(s) | Correlation with lithium use |  | Overlap with DEGs |  |
| --- | --- | --- | --- | --- | --- | --- |
|  |  |  | *r* | *P* | N genes | *P* |
| M1 | 2,092 | Transmembrane, GPI anchor, immunoglobulin | 0.156 | 9.40E-04 | 431 | 2.03E-97 |
| M7 | 700 | Helicase activity, ATP binding, metabolism, DNA replication, endoplasmic reticulum, proteasome, protein biosynthesis | -0.165 | 4.50E-04 | 22 | 1.00 |
| M9 | 55 | G-protein coupled receptor | 0.153 | 1.15E-03 | 17 | 6.15E-07 |
| M11 | 622 | - | 0.17 | 3.12E-04 | 102 | 4.93E-13 |
| M26 | 484 | Nucleic acid binding, splicing | -0.175 | 2.00E-04 | 17 | 1.00 |

Module preservation analysis was also performed to assess differences in network density and connectivity between groups, but showed full preservation indicating that networks constructed in separate groups maintain their underlying structure (Supplementary Methods and Figure S4).

### Estimated neutrophil proportions are increased in lithium users

We then sought to determine if variation in our sample could be explained by differences in blood cell-type composition. To deconvolve cellular heterogeneity, we applied CIBERSORT^33^ to our gene expression quantifications using a reference panel of 22 blood cell-type signatures. The resulting estimated cell-type proportions (Figure 2A) were then examined for their relationship with lithium use in BD cases only. Each cell type was residualized for demographic and technical variables then used to predict lithium use in a stepwise linear model. Neutrophils are the one cell type that significantly predicted lithium use within the BD cases (*β* = 0.63, P = 0.024), with elevated proportions in individuals being treated with lithium (Figure 2B). Indeed, 16 of 60 signature neutrophil genes were also lithium-use DEGs (hypergeometric OR = 4.64, *P* = 4.45 x 10^-6^).

**Figure 2.**
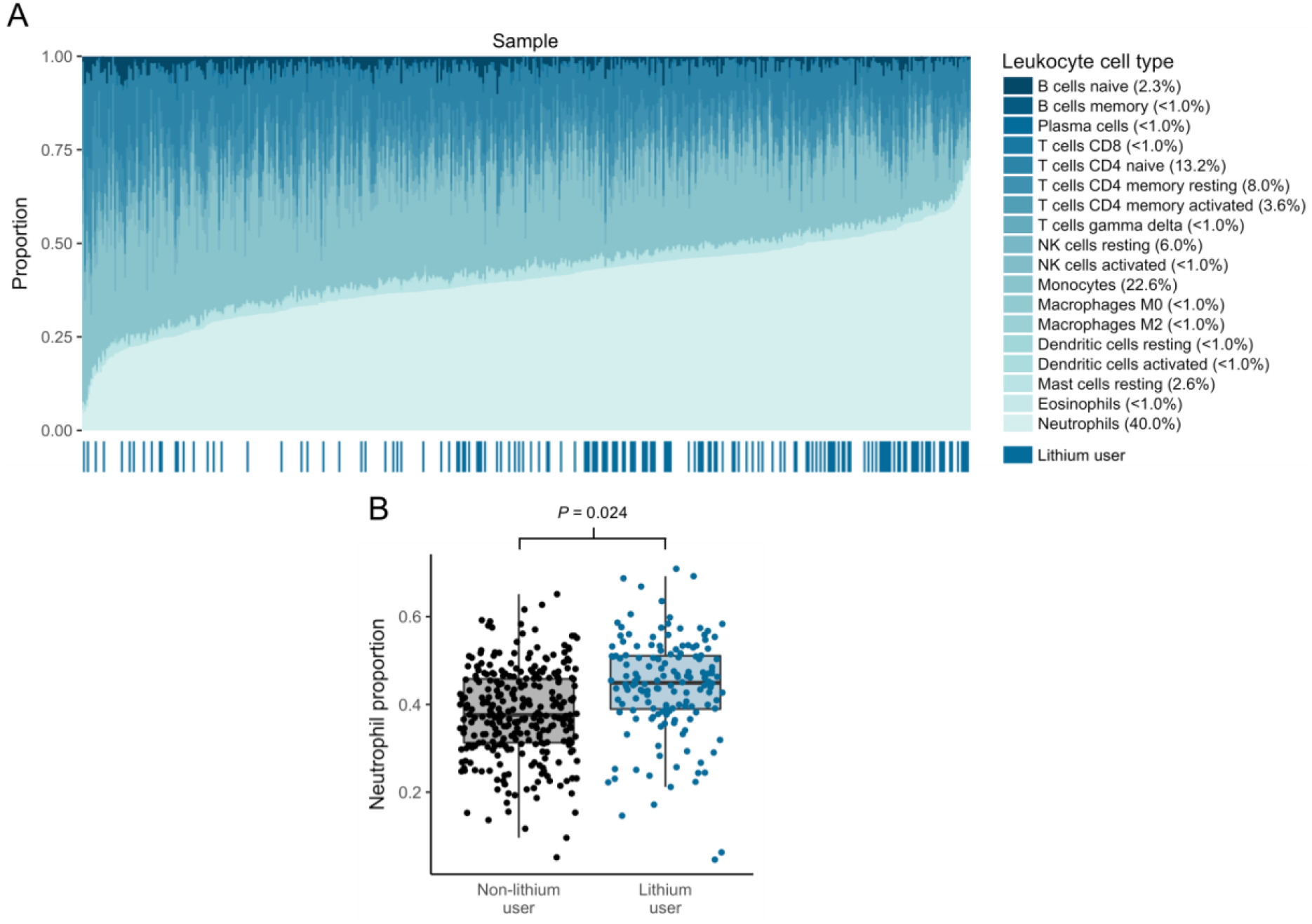
Estimated neutrophil composition association with lithium use. (A) Leukocyte cell-type proportions per sample as estimated from gene expression, sorted by neutrophil proportions. Mean proportion across samples shown in parentheses. Lithium users, shown in the bar on the bottom, cluster on the right where neutrophil proportions are higher. (B) Lithium users have higher estimated neutrophil proportions (*β* = 0.63, *P* = 0.024).

The number of genes showing differential expression in subjects undergoing lithium treatment decreased from 976 in the model without cell-type estimates to 233 in the model with cell-type estimates (FDR < 0.05; Figure S5A, Supplementary File 1), of which 194 (83.2%) were significant in the original model and concordant in direction of effect (Figure S5B). No functional annotation cluster terms remained significant after correcting for multiple testing. The number of genes differentially expressed between BD cases and controls decreased to zero after accounting for estimated cell-type proportions.

### Lithium-associated co-expression module M1 is enriched for neutrophil gene expression signatures

We then sought to determine if the various lithium-associated modules of co-expressed genes reflected biologic signatures of distinct populations of blood cell types. We did this in two ways. First, a hypergeometric overlap between lithium-associated module gene lists and cell-type signature gene lists revealed a significant overlap between module M1 with monocyte and neutrophil signature genes and M9 with eosinophil and activated mast cell signature genes (Figure 3A, left). Second, the expression of cell-type signature genes was used to predict module membership values in a linear model for each of the five lithium-associated modules. Neutrophils, monocytes, and eosinophils were again implicated (Figure 3A, right). In both of these analyses, the most significant cell type-module relationship was M1 with neutrophil estimates (hypergeometric *P* = 5.68 x 10^-21^, linear model *P* < 2.20 x 10^-16^). Indeed, neutrophil signature genes had higher M1 membership values (Figure 3B).

**Figure 3.**
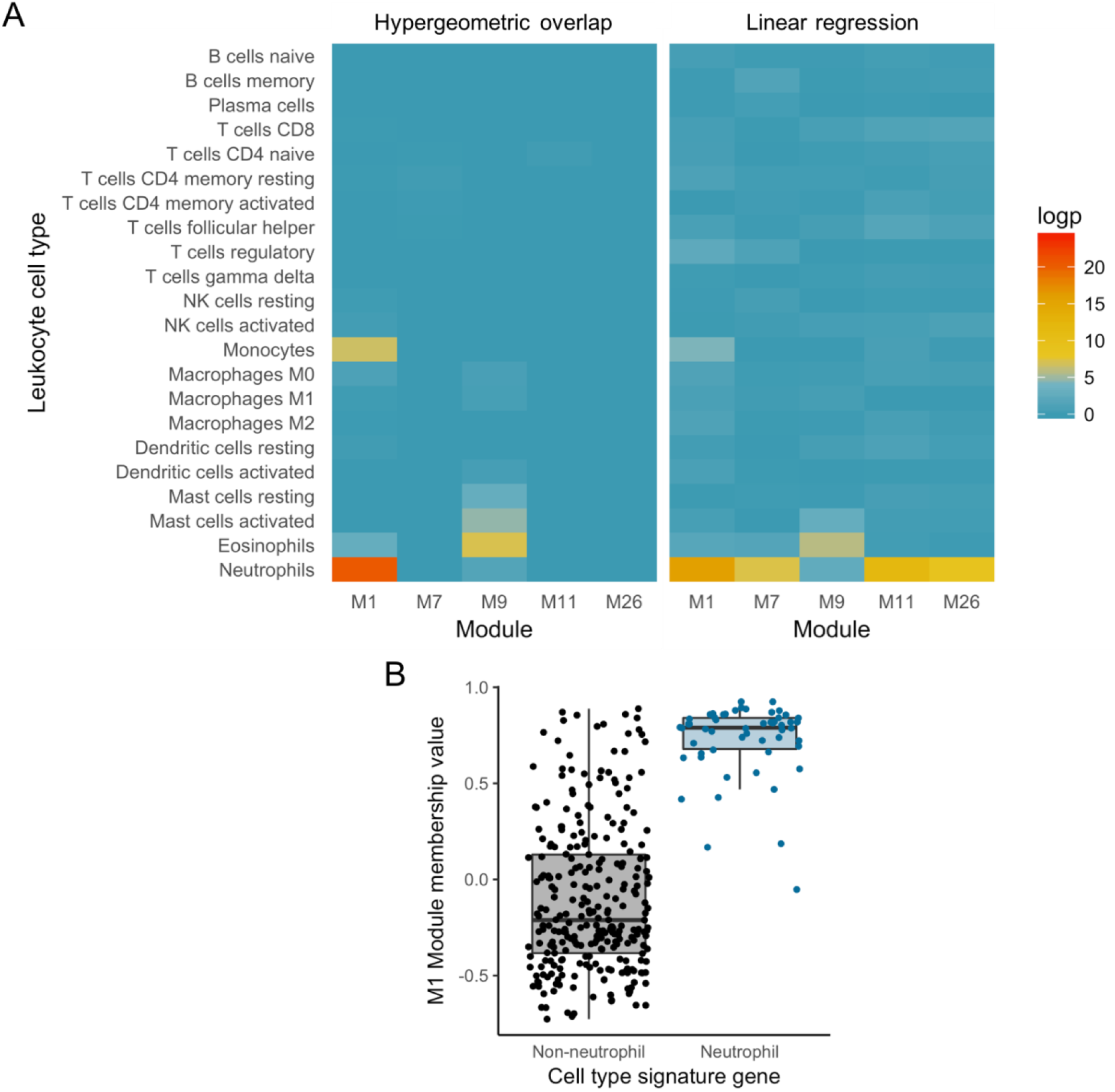
Lithium-associated co-expression module M1 enrichment for neutrophil gene expression signatures. (A) Lithium-associated module enrichment for leukocyte cell types. Left, Hypergeometric overlap between leukocyte cell type signature genes and genes in each module. Right, Linear regression of leukocyte cell type signature genes to predict module membership values. (B) Neutrophil signature genes have higher module membership values for M1 than other leukocyte signature genes (*β* = 0.60, *P* < 2.20 x 10^-16^).

### Genes with altered expression are not enriched for genes with common psychiatric risk alleles

To evaluate if BD and lithium-use DEG sets were associated with a higher burden of psychiatric risk alleles, we performed gene-set analyses using MAGMA^34^. Analyses were performed across three psychiatric GWAS traits: BD^36^, SCZ^35^, and self-reported depression^37^. SCZ and depression were used because of their high degree of overlap in SNP-based heritability with BD^4^ (Table S6). The 23andMe self-reported depression GWAS was used instead of MDD GWAS because of the large sample size and successful findings of this study. A lithium-response GWAS was not used because the SNP-based heritability estimate for this trait is not different from zero (personal communication with Thomas G. Schulze). Because the set of BD DEGs at FDR < 0.05 was too small to test, we used a more lenient significance threshold of FDR < 0.2 for this analysis instead. None of the comparisons demonstrated an association with genetic risk across the genes identified in the current study (except for the positive control gene set), even after stratifying by up- and down-regulated genes (Figure 4, Table S7). Because sleep disturbances are a hallmark of BD^39^, and due to the genetic correlation of sleep-related phenotypes with BD^40^, we performed a secondary gene-set analysis with genes implicated from chronotype, sleep duration, oversleeping, and undersleeping GWAS, which failed to demonstrate association with genes identified in the current study (Table S8).

**Figure 4.**
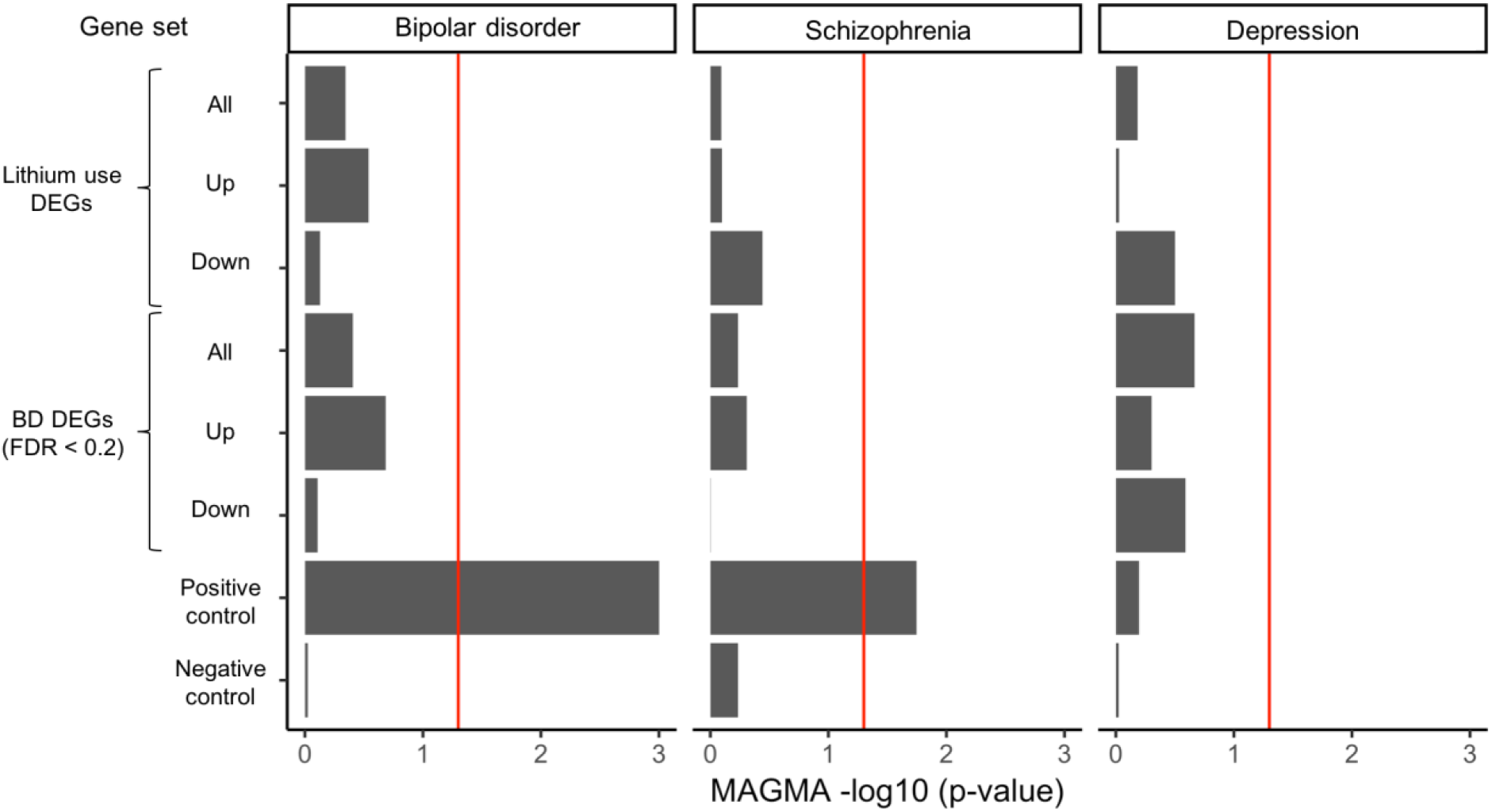
Gene-set enrichment of DEG sets with genes in psychiatric trait-associated loci (PGC BD GWAS^36^, PGC schizophrenia GWAS^35^, and 23andMe self-reported depression GWAS^37^) using MAGMA^34^. DEG sets stratified by up- and down-regulated genes. The BD DEG set was extended to include genes with FDR-corrected *P* < 0.2. The positive control gene-set consisted of the top 100 most significant genes from a random draw of N = 1,000 using the BD GWAS gene-level test statistics. The positive control gene-set association with BD was highly significant (P = 1.28 x 10^-27^) but the −log_10_ *P*-value was limited to 3 in the plot. The negative control gene-set consisted of a random draw of N = 1,000 genes using the BD GWAS gene-level test-statistics. The red line represents the significance threshold of −log_10_(0.05). All P-values and effect sizes are reported in Table S7.

## Discussion

In our whole blood BD case-control gene expression study we observed widespread subtle changes in gene expression in subjects undergoing lithium treatment but few transcriptomic differences linked to disease status. These effects were partially driven by variation in leukocyte cell type composition, and we find no evidence for a link with genetic risk for BD. Upon validation of our findings with previous *in vivo* and *in vitro* lithium treatment gene expression studies, we present a high-confidence list of genes that display altered expression associated with lithium treatment.

One of the top differentially expressed genes associated with BD, *COG4*, encodes a part of a multiprotein complex that is a key determinant of Golgi apparatus structure and capacity for intracellular transport and glycoprotein modification^41^. *COG4* mRNA is expressed widely across body tissues including the brain^42^. It has been reported as having alternative splicing in subjects with BD^43^, and concordant with our results, was reported as down-regulated in three of the ten Stanley Genomics BD brain datasets^44^. Further work is needed to determine the role of *COG4* in BD, but perhaps neuronal hyperexcitability in BD^16^ destabilizes internal cellular processes including Golgi function^45^. There is currently no evidence for a genetic link between BD disease susceptibility and *COG4^36^.*

Lithium is the first-line treatment for BD, not only for the treatment of acute episodes but also for maintenance and suicide prevention^46–47^. However, only about 30% of BD patients fully respond to lithium, it has several adverse side effects, and its mechanisms of action are not well understood^48–50^. One probable reason for this lack of understanding is the magnitude of lithium’s physiological interactions^51^. In pharmacological terms, lithium is a small molecule (the third smallest element in fact) without a defined target^50^. This lack of specificity makes it difficult to discern therapeutic mechanisms from off-target effects, which likely lead to many of lithium’s undesirable side effects and even its toxicity at doses that are too high. Lithium ions (Li+) have a single positive charge and are hypothesized to mimic and disrupt the actions and targets of more ubiquitous metal ions such as magnesium (Mg^2^+)^50^. Theorized therapeutic mechanisms of lithium include its inhibition of the protein GS3Kβ, and its effect on intracellular signaling cascades such as those involving protein kinases and phosphatidylinositol^52, 53^. It is not clear how these mechanisms relate to higher order properties thought to be involved in BD etiology like neuronal function, chronobiology, and brain structure. Examining lithium mechanisms at high biological resolution is therefore not only crucial for understanding the high rates of non-response and non-adherence to prophylactic lithium treatment in BD patients but also for understanding BD etiology itself.

The widespread but subtle gene expression changes observed in lithium users are in line with lithium’s broad scope of physiological effects^51^ and with the complex genetic architecture of BD^20^. These genes were enriched for functional annotations related to transmembrane, cell signaling, protein kinase, and immunity. These pathways have been implicated in previous BD transcriptome studies^13,14,16,23,26^ and are known targets of lithium^48, 54^. The elevated levels of neutrophil proportions we observed is in line with lithium-induced neutrophilia, which has been described since the medication’s early use in psychiatry^54^. Lithium is thought to induce neutrophilia through a complex pathway involving GSK3 and immune-related transcription factors and genes^55^. Increased levels of neutrophils are typically associated with anti-inflammatory or infection-fighting immune responses^56^. Whether these immunity-related mechanisms play a role in the mood stabilizing effects of lithium remains to be determined. Immune components of psychiatric illness including BD^57^ have long been recognized, but it remains unclear if they represent a causal pathway, a property of the disease state, or a consequence of environmental factors like body mass index or smoking. These results contribute to the understanding of the genomics of lithium action, which may be essential for the future of personalized psychiatric medicine for patients with BD. Future studies with larger sample sizes and independent replication datasets will be needed to confirm our findings, and whether these genes and pathways play a role in the mood-stabilizing mechanisms of lithium remains to be determined.

The lack of enrichment of genetic signal from common alleles associated with BD, schizophrenia, or self-reported depression suggests that genes transcriptionally associated with lithium treatment in peripheral blood most likely represent secondary effects of treatment that are independent from disease susceptibility. The lack of genetic enrichment could also indicate that our gene expression study is underpowered for this purpose, or that the transcriptomic mechanisms of genetic risk for BD are not present in whole blood. In addition, the currently available GWAS may still be underpowered thereby impacting our ability to detect a significant enrichment. With the expected rapidly increasing sample sizes of these GWAS studies we will be able to test this hypothesis more fully in the near future. We did explore the opportunity to examine enrichment of genetic susceptibility of lithium response, but because this phenotype has a SNP-based heritability not different from zero, this specific analysis is not meaningful. In this regard, it is important to distinguish between lithium *use*, the phenotype we used in our study, and lithium *response.* Self-reported answers to a lithium questionnaire by participants in our study show that the majority of subjects being treated with lithium had a positive response to the treatment and the majority of non-users have been treated with lithium in the past (Supplementary Methods). We therefore consider that the lithium use phenotype partially captures lithium response, but disentangling the complex interplay between these phenotypes is an avenue for further exploration.

Lithium use, as a trait only present in BD subjects and therefore confounded with BD diagnosis, serves as a confounder by indication and likely eliminated most of the observable BD effects. Our results highlight the importance of correcting for cell type composition as well as medication use in BD transcriptome studies. A lithium-naive study design is warranted to optimize BD transcriptomic signal that is independent of lithium use. Nevertheless, investigating the BD transcriptome in whole blood remains valuable for the following reasons. It is an accessible tissue, it has the potential for biomarker discovery, and it can be used in longitudinal study designs, which are appealing due to the episodic nature of BD. It may also be a choice tissue to observe the suggested immune component of BD etiology. In addition, peripheral tissues such as blood partially recapitulate gene expression signatures of the brain^58^, and compared to post-mortem tissues are less subject to poor quality due to rapid degradation upon death^59^. However, studies involving post-mortem tissue, *in vitro* neuronal cells, or animal models will still be needed to determine the therapeutic effect of lithium on BD-associated brain-related function.

In summary, our findings suggest that there are minimal bipolar disorder-associated gene expression changes in whole blood independent of medication use and underline the importance of accounting for such confounders in psychiatric genomic studies. While limited in their ability to uncover mechanisms associated with genetic risk, blood-based transcriptome analyses of BD may still be informative with larger sample sizes and careful designs. Lastly, our findings provide molecular insights into the potential therapeutic actions of lithium, including cell signaling and immunity-related functions. Overall, this work contributes to the understanding of BD etiology and the elusive mechanisms of its most common treatment, lithium.

## Data Availability

Gene expression data will be made available upon publication.

## Acknowledgements

We thank the study subjects for their willingness to provide specimens and clinical data. We thank GWAS research participants and researchers involved in making summary statistics available, including 23andMe research participants and employees, for making this work possible. We thank Drs. Francis McMahon and Thomas Schulze for correspondence regarding lithium-response GWAS. This work was supported by National Institute of Health grants R01MH090553 (RAO), R01MH114152 (RAO), 25602-NARSAD Distinguished Investigator Award (RAO), K99MH116115 (LMOL), and T32HG002536 (CEK).

